# Regional reinfection by Dengue: a network approach using data from Mexico

**DOI:** 10.1101/475137

**Authors:** Mayra Núñez-López, Luis Alarcón-Ramos, Jorge X. Velasco-Hernández

**Affiliations:** Departamento de Matemáticas Aplicadas, ITAM Río Hondo 1, CDMX 01080, México; Departamento de Matemáticas Aplicadas y Sistemas, UAM-Cuajimalpa, Av.Vasco de Quiroga 4871, CDMX 05300, México; Instituto de Matemáticas, UNAM-Juriquilla, Boulevard Juriquilla No. 3001, Querétaro, 76230, México

**Keywords:** Dengue dynamics, reinfection, inoculum size, human mobility, metapopulation model

## Abstract

Most of the recent epidemic outbreaks in the world have a strong immigration component as a trigger rather than the dynamics implied by the basic reproduction number. In this work we present and discuss an approach to the problem of pathogen reinfections in a given area that associates people mobility and transmission of dengue, using a Markov-chain Susceptible-Infected-Susceptible (SIS) metapopulation model over a network. Our model postulates a parameter that we have named the effective inoculum size which represents a local measure of the population size of infected hosts that arrive at a given location as a function of population size, current incidence at neighboring locations and the connectivity of the patches. This parameter can be interpreted as an indicator of outbreak risk of any location. Our model also incorporates climate variability represented by an index based upon precipitation data. We replicate observed patterns of incidence at a regional scale using data from epidemics in Mexico.

## I. INTRODUCTION

It is well known that the global spread of a disease strongly depends on the number of secondary cases generated by a primary case in susceptible populations and also on the time it takes to infect those secondary cases (see, for example, [36]). Transportation and movement of human populations and of vectors of disease contributes to the spread by allowing contact between new susceptible populations both in vectors and human hosts that may be regionally or geographically separated from the focal point of the first outbreak. This interplay has been addressed from a variety of perspectives in the literature but we are particularly concerned in this work with the spread of vector-transmitted diseases (see [10] and references there in). It is well known that connectivity between population centers and travel are closely related to the import/export of infectious diseases both in directly-as well as in vector-transmitted diseases and that patterns of human mobility are seasonal [40]. Climatic conditions affect transmission since pathogens life cycles and habitat suitability for vectors, hosts or pathogens can be significatively modified by it. Human mobility has also a seasonal component [40]. Dengue fever is a re-emerging mosquito-borne infectious disease that is of increasing concern as human travel, migratory patterns and expanding mosquito ranges increase the risk of spread [18], [30], [27], [39], [40]. Dengue has caused illness in millions of people over the last several years [41] and concerns have been expressed about the increased risk it posses on the general population due to the impact of climate change [33]. Dengue is only one of the vector-born disease that has had major outbreaks in several countries around the globe. Recently, Chikungunya virus caused outbreaks in many Caribbean [21] and several Indian Ocean islands [2] and Latinamerica including Mexico [3]. Zika, another vector-borne disease transmitted by the same mosquito species (*Aedes aegypti*), had a major epidemic in the American continent in the year 2016 and its introduction into anon-endemic zone (Florida) has been addressed form the modeling point of view in [8].

Modeling and analysis of the spread of infectious diseases has become a relevant problem of interdisciplinary nature. Many countries have established public health surveillance systems geared to the prevention and rapid response in case of an epidemic outbreak and mathematical and statistical models are being amply used as tools in public health [17], [18].

Dengue modelling and control efforts are extensive [17], [26], [34], [37], although there is much still to do. The primary vector for dengue is *Aedes aegypti* and this virus generates acute immunizing infections in humans. Dengue is primarily transmitted by *A. aegypti*, but *Ae. albopictus* can be an important secondary vector. Both mosquito species are diurnal, biting mostly in the morning and evening rather than at night [28]. Dengue is in itself a complex disease with multiplicity of lineages [4] that causes a spectrum of illnesses in humans ranging from clinically inapparent to severe and fatal hemorrhagic disease. Classical dengue fever is generally observed in older children and adults and is characterized by sudden onset of fever, frontal headache, nausea, vomiting and other symptoms [35].

Mitigation strategies for dengue include reduction of the mosquito population via indoors praying (adulticides) larvicides, lethal ovitraps, removing man-made oviposition sites and reduction of human exposure to mosquito bites via the use of screens, mosquito repellent, etc., but these are not always effective, as a consequence, the absolute numbers of Dengue infection have increased during the last 40 years [12]. Unfortunately, countries where positive results exist for the vector eradication have been suffering from epidemics outbreaks: the disease is coming back. Vaccines are also in development and currently in clinical trials [14], [17].

Human movements result in global spread of infectious diseases [31], [39], including vector-borne diseases [19]. Regarding to the role of mobility in the dengue dispersion, Martínez-Vega *et.al.* [30], have demonstrated that the spread of infection within a locality mainly depends on human mobility. For example, subjects between 30 and 64 years old, despite they probably have lower force of infection, being asymptomatic and economically active, move to daily destinations where they remain long enough to be bitten by nearby vectors, transmitting dengue to other close subjects which go to their homes and start a new peridomestic transmission cluster. Young and elderly subjects would have a lower participation in the local dengue dissemination because they have limited mobility, since younger individuals are typically symptomatic while older ones have lower mobility. In [25], the risk of infection importation and exportation by travellers is estimated, taking into account the force of infection of the disease in the endemic country to disease-free countries. A similar situaton has been addressed in [8] where the importation of cases is explicitly modeled.

In recent years, spreading processes, i.e. computer viruses in networks, epidemics in human populations, rumors or information in social networks, are modeled and described by complex networks. Each node of the network represents an element of the system and the links represent the interaction among nodes. Recently, Markov-chain based models for a Susceptible-Infected-Susceptible (SIS) dynamics over complex networks have been used to describe the dynamics of individual nodes and to determine macroscopic properties of the system [6], [7], [15], [16],[31], [38]. An important result from Markov-chain based models, is the presence of an infection threshold that depends on the value of the spectral radius of the adjacency matrix.

In this work, inspired by ideas of [6], [7], [38], we propose and discuss a model that associates people mobility and the transmission of dengue using a Markov-chain based model for a Susceptible-Infected-Susceptible (SIS) dynamics over complex networks. It is known [8] that immigration processes in deterministic models render *R*_0_, the basic reproductive number, ineffective as a threshold and invasion parameter and the model analysis becomes substantially more difficult compared to the no-immigration (standard) case. We attempt in this work to overcome this problem and address specifically the reinfection process of whole geographical regions and only indirectly we follow particular infections of individuals. We have chosen a patch-dynamics approach to the Dengue colonization-extinction processes of sites. In a network of sites, each site can be reinfected (recolonized) by the movement of infectious individuals from neighboring patches and the disease in any given patch may decline due to the natural disease life cycle or because of emigration of sick individuals. Each node in the network that we define in this work corresponds to populations in four different states of Mexico: Veracruz, Guerrero, Oaxaca and Chiapas. The links in the network and the interaction among the nodes, are modeled according to their shared borders as will be described in the following section. Additionally, the model incorporates external forcing as a way to explore the effects of climate variability on the spread of dengue, particularly rain. Our model attempts to predict the probability of occurrence of an outbreak (colonization-invasion) in any node in the network. The parameters of the model are fitted to real data and simulations are performed to show and compare the model behavior with the observed patterns of incidence. The paper is organized as follows. In Section 2 we present the parameter estimation and discrete mathematical model for dengue spread in a large geographical region. In Section 3 we present and discuss the numerical results of the outbreaks. Finally, in Section 4 we draw some conclusions about this work.

## II. THE MODEL

The classical Levins metapopulation model [22] has been generalized to many contexts, in particular to spatially explicit settings where it has shown that landscape structure and patch dynamics can alter the dynamics and persistence of metapopulations (for example [20]). Landscape structure, characterized by spatial and temporal heterogeneities determines the nature and impact of all of these effects [20]. Here, we formulate a mathematical model following the classic work of Levins [22], [23] in an spatially-explicit version. It is a patch dynamics model with time-dependent contact (infection) rates associated to climatic variability, particularly rain. We are interested in describing the population dynamics of the dengue infection reported for the years 2004 to 2009 in Mexico as the infection moves through four patches where migration rates depend on the existence of a common border between neighboring states. It is well know that for Dengue, population mobility is a very important factor for transmission. In Mexico population mobility is essentially carried out on buses through the road network of the country. We concentrate in a relatively simple system constituted by large regions, in this case political subdivisions known as states. Our data base comprises weekly incidence data reported to regional health centers in each of four states whose names are Oaxaca, Guerrero, Veracruz and Chiapas (see map Fig 1).

**FIG. 1:**
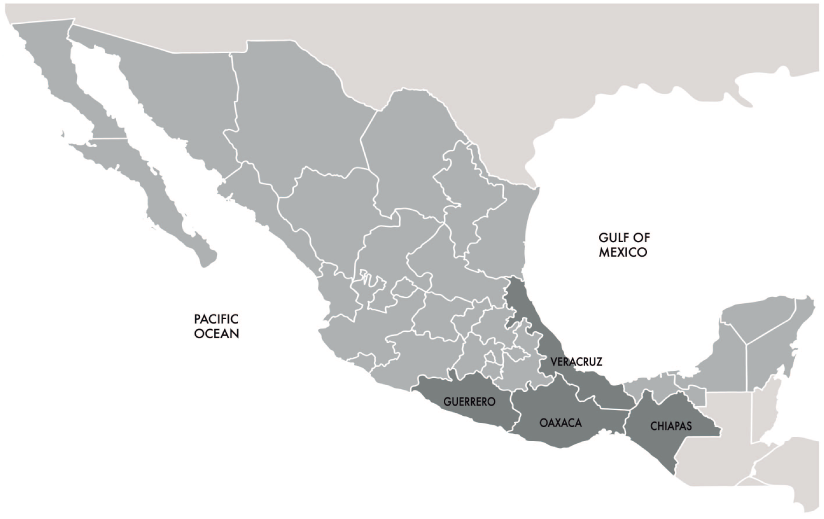
Data shows weekly incidence data from the National System of Epidemiological Surveillance, of four states indicated by the darker color in the map: Oaxaca, Guerrero, Veracruz and Chiapas.

The information has been aggregated by state. There are certainly alternatives to the choice of spatial aggregation done here. For example, we could have chosen to identify regions by hydrological basins since this subdivision could be associated to mosquito data in a more natural way. However, lacking information on vector abundance and distribution our choice was to look at aggregated data following state boundaries. However, we are currently preparing a forthcoming work where some of us we look at hydrological basins and roads associated to epidemic outbreaks.

### A. Model parametrization

Our information from the National System of Epidemiological Surveillance covers the history of epidemic events for several years in different municipalities of South Eastern Mexico. The recorded cases are provided by Regional Hospitals that are in charge of reporting infectious diseases to the health authorities in the country. Dengue is a mandatory notifiable disease in Mexico but only diagnosed cases are reported so we can expect our data to be biased by un-notified asymptomatic cases, under-reporting and misreported (a certain amount of cases that are not related to Dengue but to some other diseases with similar symptoms). There is evidence in the Dengue literature [13], [42], that movement is a key factor for understanding epidemic episodes and large outbreaks that occur along the years. Here we analyze data corresponding to towns located in four states in the Federal Republic of Mexico: Veracruz, Guerrero, Oaxaca and Chiapas. The source of information that we have are the time series of cases from the years 2004 to 2009. The time series for Dengue cases is presented in Fig. 2.

**FIG. 2:**
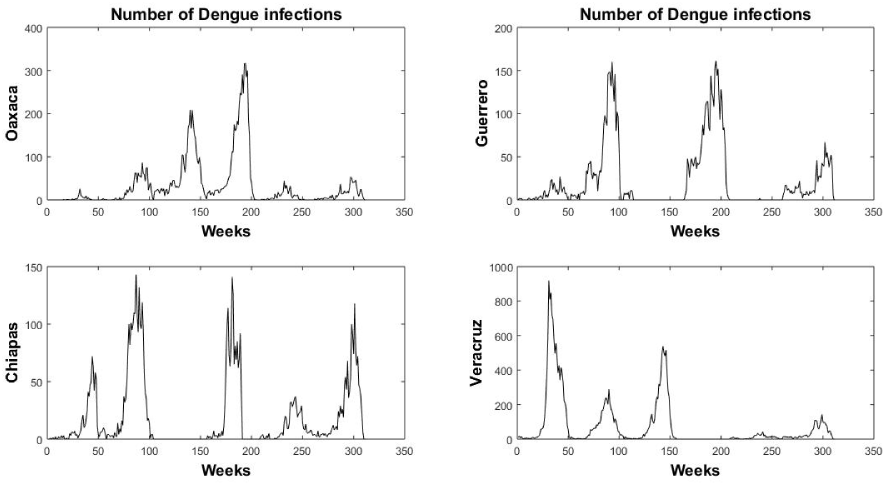
Weekly number of dengue cases aggregated by state reported in South Eastern Mexico: Oaxaca, Guerrero, Chiapas and Veracruz (2004-2009).

To be able to parametrize our model we need an estimation of the force of infection or of the infection rate for each outbreak which we obtain through the estimation of the basic reproduction number. Once the estimate of each reproduction number is obtained, we deduce the infection rate for each State used to generate our simulations.

As it is well known, the reproduction number, *R*_0_, is the expected number of secondary cases produced in a susceptible population by a typical infective individual during the time in which s/he is infectious. If *R*_0_ < 1, then on average an infected individual produces less than one new infected individual over the course of its infectious period and the infection cannot grow. Conversely if *R*_0_ > 1 then each infected individual produces, on average more than one new infection and the disease can invade the population. From its definition *R*_0_ is determined from early stages of the epidemic and its magnitude is a useful indicator of both the risk of an epidemic and the effort required to control an infection. We could have used a better estimate, for example the effective reproduction number developed in our group [1], but the data base has many weeks in which the number of cases is zero. This fact introduced technical difficulties and inaccuracies in the estimation by that method and after consideration, we decided to discard since the methodology described in the following paragraph rendered better and more robust results.

#### 1. Exponential growth rate method (EG)

The rate of exponential growth *r* is defined as the per capita change in number of new cases per unit of time (epidemic growth rate). One of the ways of inferring *R*_0_ from *r* is through a moment generating function expression for the reproductive number. According to [43], the reproductive number defined as expected secondary infections is given by 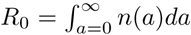, where *a* is the time since infection, *n*(*a*) expected rate of generation of secondary cases at time *a* since infection. The rate *n*(*a*) can be normalized to a distribution *g*(*a*), *i.e. g*(*a*) = *n*(*a*)*/R*_0_. The Lotka-Euler equation is given by

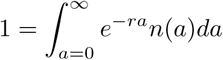

and substituting *g*(*a*) (generation interval distribution) into it we obtain

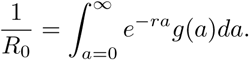

The term in the right-hand side, is the moment generating function *M* (*z*) of the distribution *g*(*a*), *i.e.* 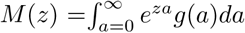 [44]. Evaluating at *z* = −*r* we obtain the reproductive number as 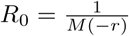.

During the early phase of an outbreak, the exponential growth rate is defined by the per capita change in the number of new cases of infected people per unit of time. To estimate *R*_0_ from the real data we choose a period in the epidemic curve over which the incidence growth is exponential and then a Poisson regression to estimate it (rather than linear regression of the logged incidence [32]). As an example, in Figure 3, an initial inspection of the incidence data shows that the exponential growth period occurs during the first 23 weeks for Oaxaca during the year 2005. Important parameters in an epidemiological model are the per capita contact rate between susceptible and infected individuals *β* and the infection recovery rate *µ*. No births or deaths are taken into account given the time frame of an individual epidemic outbreak.

**FIG. 3:**
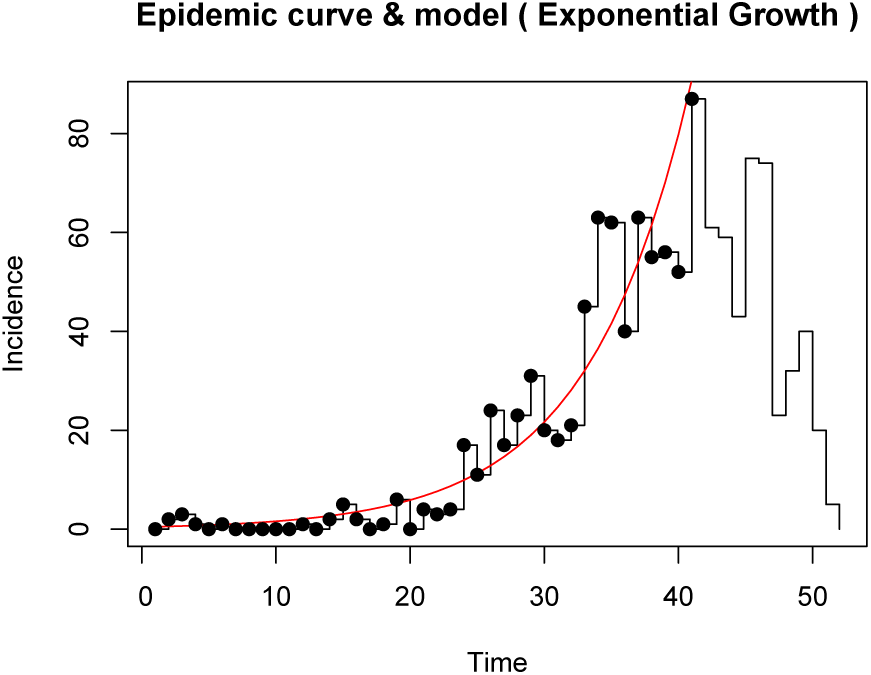
Observed incidence and predicted incidence for Oaxaca State during 2005.

The incidence data is given as number of new cases per epidemiological week and we use a time dependent maximum likelihood method with the distribution of generation times rescaled to weeks to obtain the values reported in Table 1. This Table lists the estimated values of *R*_0_ for each year for Oaxaca State. Once the estimate of each reproduction number is obtained, we solve for the infection rate *β* [45]. In the Appendix all the *R*_0_ tables for the rest of the States are displayed.

**TABLE I:**
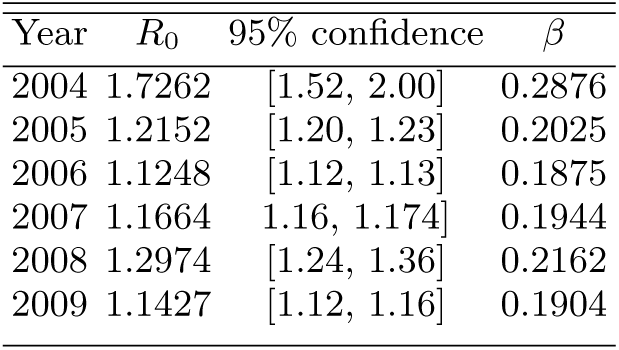
R_0_ for Oaxaca State estimated with the exponential growth rate method with a mean infection period of 6 days.

| Year | $R_0$ | 95% confidence | $\beta$ |
| --- | --- | --- | --- |
| 2004 | 1.7262 | [1.52, 2.00] | 0.2876 |
| 2005 | 1.2152 | [1.20, 1.23] | 0.2025 |
| 2006 | 1.1248 | [1.12, 1.13] | 0.1875 |
| 2007 | 1.1664 | 1.16, 1.174] | 0.1944 |
| 2008 | 1.2974 | [1.24, 1.36] | 0.2162 |
| 2009 | 1.1427 | [1.12, 1.16] | 0.1904 |

### B. Mathematical model

Following [38] we take each federal state (Oaxaca, Veracruz, Guerrero, Chiapas) as a node of the network and define that it can be in either of two (epidemiological) states in analogy to an SIS epidemic model.

The discrete-time Markov-chain spreading model for dengue transmission is set up under the following assumptions:

- Each node of the network represents the population of each one location (heretofore renamed as *locations* or *sites*) of the Mexican Republic: Chiapas, Guerrero, Oaxaca and Veracruz. The connection between locations is defined by adjacency: having a common border; see Figure 4. Each location can be in either of two epidemiological *states*: *S* susceptible (empty of infection) or *I* (occupied or infected).
- The probability of finding an *infected* individual in location *i* at time *t*, is defined as *p_i_*(*t*), with *i ∈ E* = {Chiapas, Guerrero, Oaxaca and Veracruz}. Therefore, each *location* at each time *t* is with probability 1 − *p_i_*(*t*) in *state S* (without outbreaks), and with probability *p_i_*(*t*) in *state I* (with outbreaks).
- We assume that Dengue infections are climate sensitive, due to the dependence of the mosquito life cycle on precipitation, humidity, temperature, and so forth; moreover vectorial capacity is a highly climate-sensitive parameter [24]. Dengue outbreaks vary over characteristic periods longer than a year and the climatic variability drives these cycles. In this sense, precipitation is chosen as a general “proxy” of an external factor, that may change the probability of contagion [5].

**FIG. 4:**
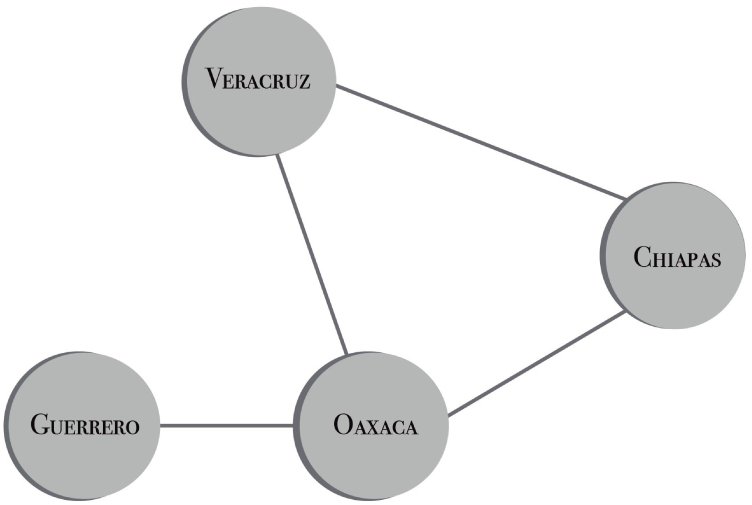
Network of locations where outbreaks are propagated. The infection moves through four patches where migration rates depend on the existence of a common boundary betweenneighboring states.

We are given a network of *N* nodes (locations) and some directed links between them. We assume discrete time-steps of size ∆*t*, corresponding to an epidemiological week. With all of the above, the transition between states (*S* and *I*) will depend on the probability of infection and the movement of infectious and susceptible individuals between the locations or nodes of the network. Figure 5 shows the transition between *state S* and *state I*. Note that each node in Figure 4 has associated the graph presented in Figure 5.

**FIG. 5:**
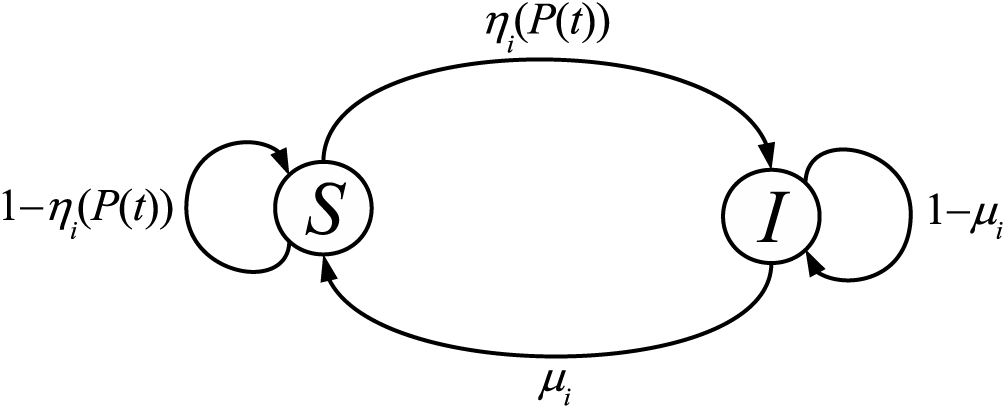
Transitions for each node showing the two possible states for each location, and the transition probabilities between states *S* and *I*.

The aim of this study is to determine η_*i*_(*P* (*t*)), which we call the *interaction* function, associated with the Dengue outbreak data shown in Fig. 2. Following [2, 3] this interaction function represents the probability that location in the network is infected or colonized by interacting with its neighbors.

Any location in state *I* recovers and passes to state *S* with probability *µ* (clearance or recovery rate), but it is reinfected with probability η_*i*_(*P* (*t*)), where

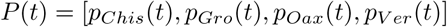

is a vector with entries *p_i_*(*t*) that represent the probability of outbreaks in location *i* at time *t*.

The probability function η_*i*_(*P* (*t*)) quantifies the probability that in *i*th location, the number of outbreaks will increase due to population displacement from the *j*thlocation with *i* = *j*; this probability is affected by climatological factors (precipitation) as will be shown later.

The function η_*i*_(*P* (*t*)) is constructed as follows: suppose that location *i* receives, on average from a neighboring location *j*, *r*_*i*_*N*_*j*_ individuals, where *N*_*j*_ represents the population in location *j* and *r*_*i*_ the fraction of individuals that moved from *j* to *i* in each step of time. In this migrant population, there are on average *r_i_N_j_p_j_*(*t*) infected individuals (on average there exist *N_j_p_j_*(*t*) Dengue cases in location *j*) with *i, j ∈ E*. Therefore, the *effective infective inoculum size* arriving to location *i* will be given as the sum of i) immigrant individuals that enter that location, plus ii) the cases that already exist at that location and iii) minus the infected individuals leaving *i* (emmigration), given by expression that enter that location, plus ii) the cases that already exist at that location and iii) minus the infected individuals leaving *i* (emmigration), given by expression

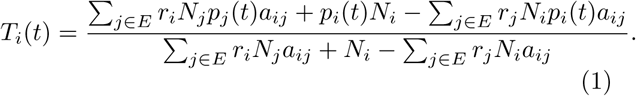

*A* = (*a*_*ij*_) is adjacency matrix of the network (see figure 4) where *a_ij_* = 1 if *i* = *j* or if locations *i* and *j*, *i ≠ j*, are neighbours, otherwise *a*_*ij*_ = 0 for *i, j ∈ E*.

Now by definition η_*i*_(*P* (*t*)) is a function of three factors: the first one is the probability β_*i*_ of an individual in location *i* becoming infected which is defined as the product of the per-contact probability of infection times the per capita number of contacts per unit time that we assume constant for each location *i*; the second factor is the effective infective inoculum size *T_i_*(*t*), and the third is the average monthly (lacking information on weekly precipitation, our choice was to consider as an approximation, the same precipitation for the weeks corresponding to the same month) precipitation in location *i* given by *f_i_*(*t*).

Precipitation data was obtained from Sistema Nacional de Información del Agua (SINA) [9], Mexico, from 2004 to 2009 for each location during the time period corresponding to the epidemic data. We have incorporated climate variability *f_i_*(*t*) represented by an index based upon monthly precipitation data, therefore all the weeks of a month have the same value.

The recovery rate given by *µ* (recovery rate), can be obtained from various studies, for example [14, 30]. In the model this parameter is constant throughout the simulation.

Our model is a discrete time Markov process dynamical system [38], described as follows

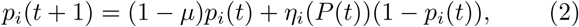

where *i ∈ E* = Chiapas, Guerrero, Oaxaca, Veracruz and 0 ≤ *p_i_*(*t*) ≤ 1. Time is discrete, each time step corresponding to an epidemiological week.

## III. RESULTS

We start first with a simple case for our probability function η. We incorporate variable contact rate but do not explicitly include the effective inoculum size *T*. This factor will be introduced in the next case. For now let

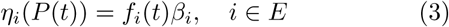

note that *f*_*i*_(*t*)β_*i*_ gives the time-dependent infection rate of location *i*, so in the discrete mathematical model simulation the forcing function with periodicity multiplies the contact rate β_*i*_.

Incidence is a term commonly used in describing disease epidemiology. Incidence is the rate of new cases of the disease. It is generally reported as the number of new cases occurring within a period of time; in this work the incidence is per week.

In Figure 6 we show the aggregated data (incidence) summing all cases in all locations of the outbreak data for the years 2004-2009 (the *x*-axis represents epidemiological weeks) compared to simulated data from equation (2) with η_*i*_(*P* (*t*)) given by Eq. (3), with the network average probability of an outbreak computed as

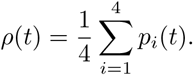

both for real and simulated data. Note that, in general, we obtain a good fit for the observed data although in the last two outbreaks their amplitude is overestimated by our model but nevertheless it accurately reproduces the timing.

**FIG. 6:**
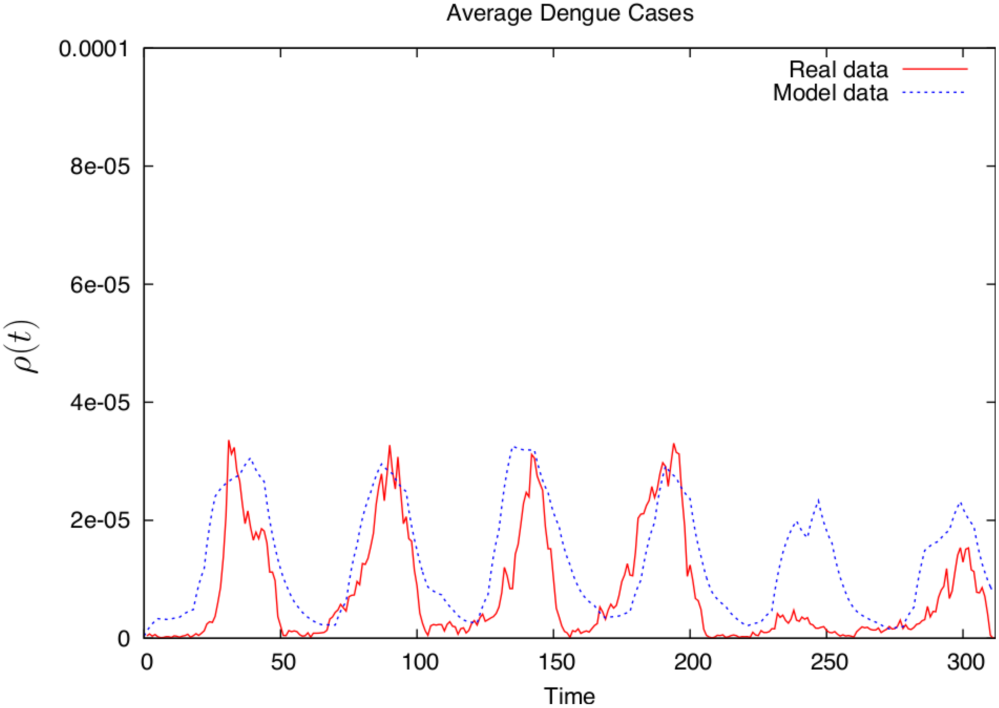
Model simulations compared with actual data for the period 2004-2009 without the incorporation of human mobility.

Now we proceed to include host mobility in order to obtain a more realistic scenario to compare with our data. To incorporate the displacement of infectious people into the probability function, what we have called before the *effective inoculum size*, we use the following explicit expression for (1):

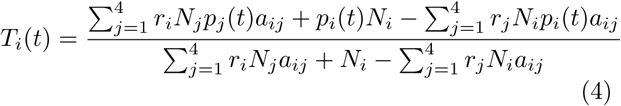

where *a*_*ij*_ is an element of the adjacency matrix that establishes the existence of a connection between locations *i* and *j* (defined as above).

According to Eq. (2), we describe the colonization-extinction of cases by counting cases moving to location *i*, cases that stay in location *i* and those that recover in location *i*:

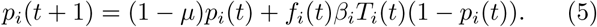

Recall that data on precipitation for each location is considered. To this effect we use the maximum precipitation of each year to normalize and define a precipitation index with maximum value of 1 (see Figure 7).

**FIG. 7:**
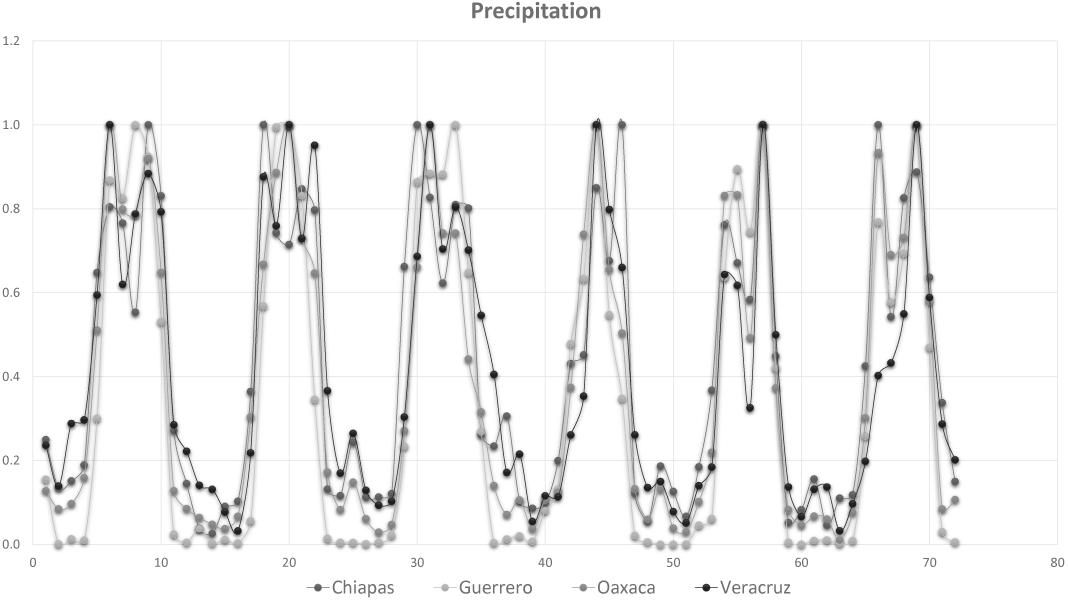
Standardized precipitation data for the period 2004-2009.

Data on mobility through public transportation in the roads and highways of Mexico is scarce and very incomplete. However, using the information on the total population in each location (federal state) and the economic strength of each of the Mexican states considered, then *r_i_*, the fraction of individuals that move to location *i*, can be roughly approximated. Here we set as a hypothesis that individuals migrate in higher proportions to Guerrero State, followed by this, the state of Veracruz, Chiapas and Oaxaca [11] the hierarchy defined from the less populated, poorer location to the most populated and better-off location.

Figure (8) shows a comparison between real and simulated aggregated outbreaks for the whole network considering the mobility corrected parameter *T_i_*(*t*). Observe that relative to the results shown in Figure (6), the fit has significantly improved.

**FIG. 8:**
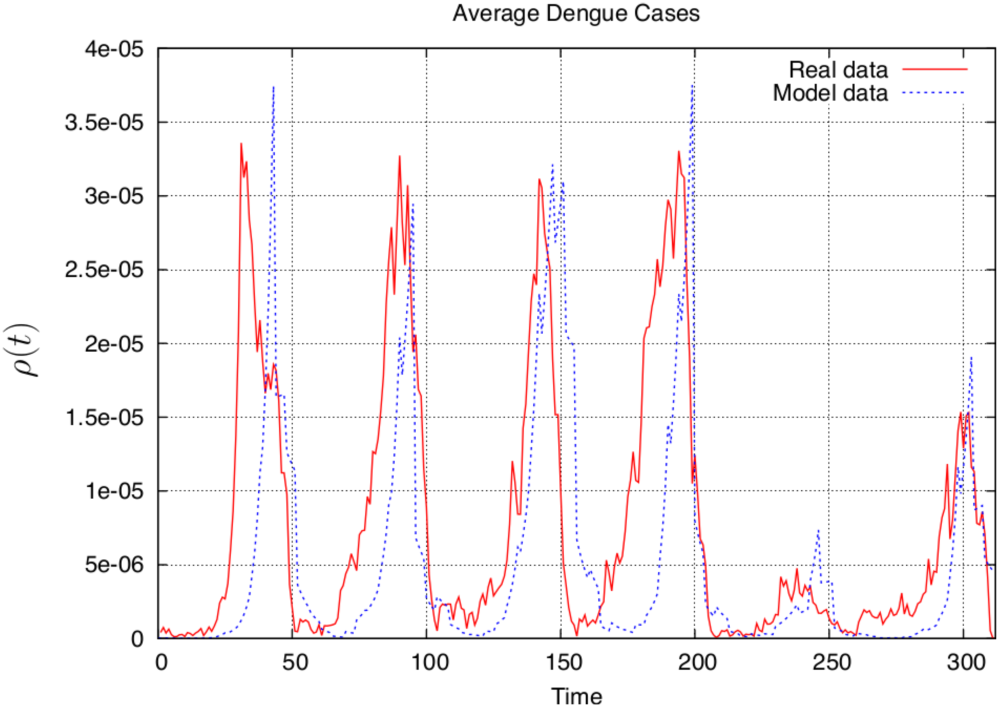
Comparison between average observed outbreaks and numerical simulations for the whole network for the years 2004-2009 incorporating host mobility (cf. Figure (6)).

Our data and simulations are scaled to proportions so our fit is qualitative in that sense. However, note that the phase of the outbreaks is slightly delayed by one or two weeks but both the amplitude and the interepidemic periods, which in general are events with very low numbers in the data, are well approximated.

In Figures (9)-(12) we show the comparison for each location (federal state) with a qualitatively reasonable approximation for Oaxaca and Veracruz states. In both cases the slight out of phase estimation is evident but once again both the amplitudes and interepidemic periods have a good qualitative approximation.

**FIG. 9:**
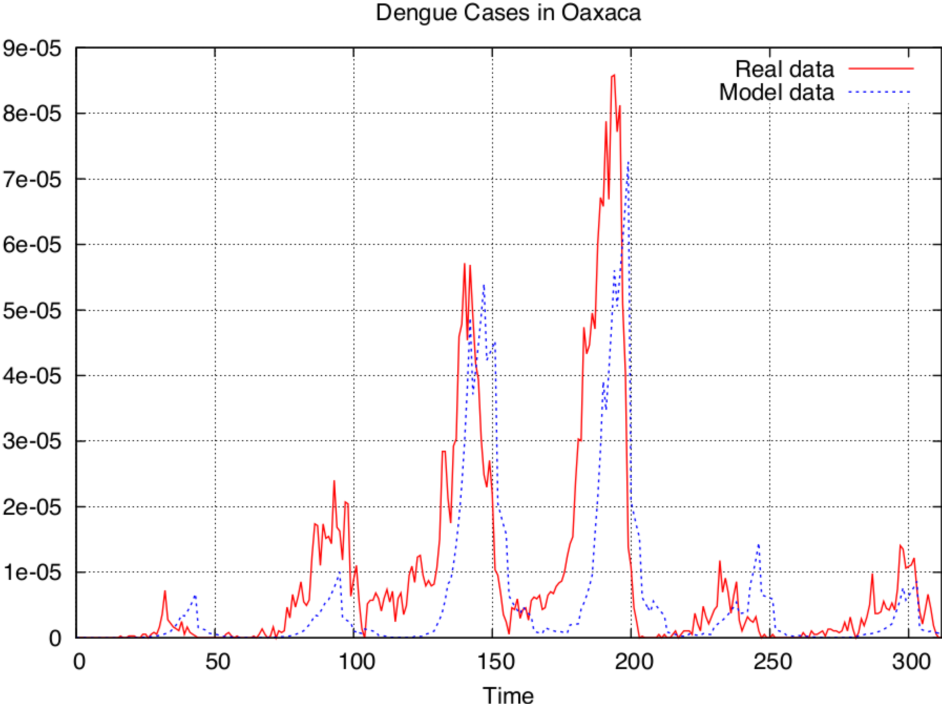
Comparison between observed outbreaks and numerical simulations for Oaxaca State during 2004-2009 with host mobility.

**FIG. 10:**
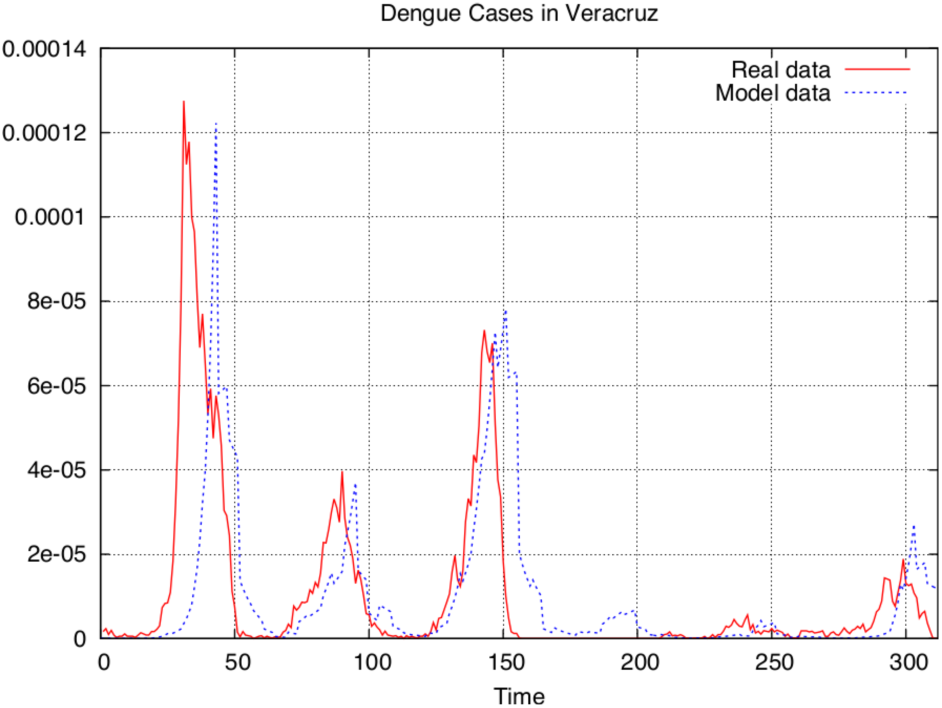
Comparison between observed outbreaks and numerical simulations for Veracruz State during 2004-2009 with host mobility.

**FIG. 11:**
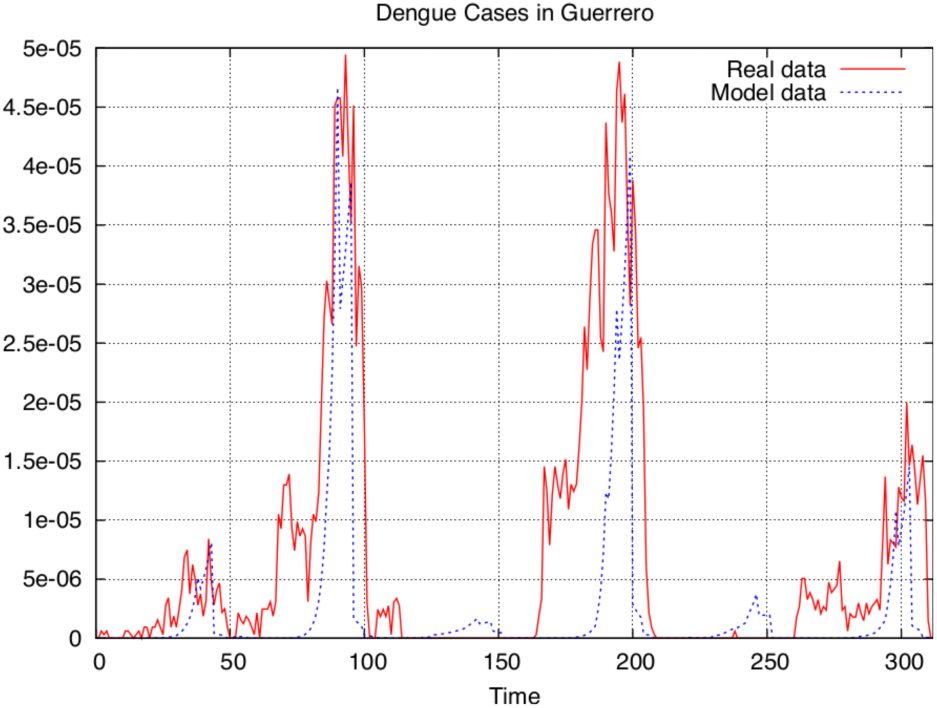
Comparison between observed outbreaks and numerical simulations for Guerrero State during 2004-2009 with host mobility.

**FIG. 12:**
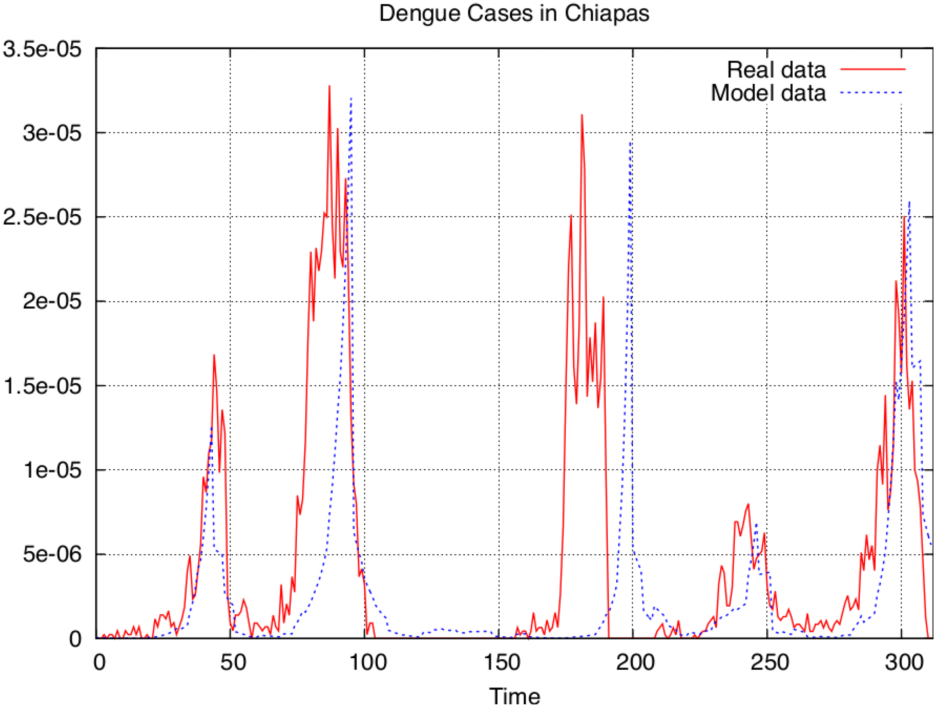
Comparison between observed outbreaks and numerical simulations for Chiapas State during 2004-2009 with host mobility.

However in Chiapas and Guerrero States, our model does not perform well. For Guerrero the phase of the outbreaks misses two outbreaks and the amplitude of the epidemic curves is overestimated. For Chiapas we obtain spurious outbreaks. It is important to mention that Guerrero and Chiapas are states with very many weeks with no reported cases and that this scarcity of cases is very likely the cause of the relatively bad fit of our model.

## IV. DISCUSSION

We have used a very classical SIS metapopulation model to approximate the processes of reinfection in large geographical areas that we view as large patches using, as a first approximation, incidence data available by location.

Given the very complex interplay of the different dengue serotypes and the lack of data pertaining to their incidence or prevalence, we have aggregated the incidence data into a process geared toward the understanding of the geographical dynamics of Dengue using a colonization-extinction framework. This framework, albeit simple in terms of the actual population dynamics of Dengue, concentrates on the movement patters underlying the spread of this disease in a large region of Southern and Eastern Mexico.

Our model is stochastic in nature and we have incorporated into it climate variability represented by an index based upon monthly precipitation data as an approximation to more general indexes of climatic variability. A new parameter is defined as *T*_*i*_(*t*) the *effective infective inoculum size* which represents a local measure of the population size of infected hosts that arrive at a given location as a function of population size, current incidence at neighboring locations and the connectivity of the patches. This parameter can also be interpreted as an indicator of outbreak risk of location *i*. From our results one can see that the location with the lowest degree (Chiapas) is the one with the poorest incidence fit. Our data lacks locations that have common boundary with the Guerrero location and, therefore, the outbreak risk index *T*_*i*_(*t*) for this location does not integrate all the real inputs and outputs acting of the site. The other nodes in the network present a better fit since incidence data from first neighbors is available.

With this very simple framework we are able to very closely reproduce the incidence dynamics in all four locations considered except for a number of outbreaks where incidence data is particularly scarce. We interpret this result simply as a verification that, regardless of the complexity of the population dynamics of Dengue, movement at a geographical scale is a relatively simple colonization extinction process taking place in a network (an spatially extended system) whose dynamics is dependent on its topological arrangement, and neighborhood interactions [22], [29]. Our model considers, as in [40], a connectivity matrix in which population flow changes as a function of climatic variability (precipitation), total population size of connected sites and weekly incidence. Our results indicate that the directional mobility of human populations in the four sites considered here, changes seasonally in a given year.

During the years covered by our data, the Dengue strains that have circulated have been mainly Dengue II and I with lower prevalence of Dengue III and IV. Immunity, therefore, must play a role in the observed reinfection dynamics. However, since we are aggregating all data at the level of the location, i.e., we are adding all cases in all the regional hospitals that report them each week, we miss, at this very large spatial scale, the impact of immunity. In our data, Dengue outbreaks at the town or municipality level are notoriously asynchronous and there are large gaps with no cases in the weekly records. We argue that this is the reason why immunity apparently imposes no important bias on the fitting of incidence that we get. To improve our colonization-extinction model, an ongoing effort is been made to generalize and adapt our results is the SIS model to the ideas of [20] where patch unsuitability is introduced into the patch classes of Levins’ model. In our case unsuitable patches for colonization would be locations with a large population of immune hosts. This approach should improve our estimates and shed light into the dynamics of movement in Dengue. The main focus of this paper has been the examination of the role of the inoculum size and precipitation effects. For future work the different patterns both geographical and temporal of precipitation and temperature that affect mosquito population size will be explored for their impact in the detail dynamics of the outbreaks.

## Acknowledgments

We thank Rogelio Danis-Lozano for providing the dengue data used in this paper. MNL acknowledges the financial support from the Asociación Mexicana de Cultura, A.C. JXVH acknowledges the support from the NoMMA (Multidisciplinary Node of Applied Mathematics) and LIIGH (International Laboratory for Research in Human Genome) at UNAM-Juriquilla and from grant PAPIIT IN110917.

## Appendix

The basic reproduction number *R*_0_ is defined as the number of secondary infections that a single infectious individual produces in a population where all hosts are susceptible.

During the early phase of an outbreak, the exponential growth rate is defined by the per capita change in number of new cases of infected people per unit of time. For the calculation of *R*_0_ from the real data is necessary to choose a period in the epidemic curve over which growth is exponential. We implemented the exponential growth method to estimate this parameter, for the purpose of calculating the infection rate *β* according to the infection period in each location.

**TABLE II:** *R*_0_ and infection rate *β* of Oaxaca State with an infection period of 6 days.

| Year | $R_0$ | 95% confidence | $\beta$ | 95% confidence |
| --- | --- | --- | --- | --- |
| 2004 | 2.9424 | [2.39, 3.71] | 0.4904 | [0.40, 0.62] |
| 2005 | 4.0434 | [3.82, 4.28] | 0.6739 | [0.64, 0.71] |
| 2006 | 5.4739 | [5.19, 5.76] | 0.9123 | [0.87, 0.96] |
| 2007 | 7.4272 | [7.20, 7.66] | 1.2378 | [1.20, 1.27] |
| 2008 | 3.0005 | [2.69, 3.34] | 0.5000 | [0.49, 0.56] |
| 2009 | 2.4799 | [2.27, 2.70] | 0.4233 | [0.39, 0.47] |

**TABLE III:** *R*_0_ and infection rate *β* of Oaxaca State with an infection period of 14 days.

| Year | $R_0$ | 95% confidence | $\beta$ | 95% confidence |
| --- | --- | --- | --- | --- |
| 2004 | 2.9424 | [2.39, 3.71] | 0.2102 | [0.17, 0.26] |
| 2005 | 4.0434 | [3.82, 4.28] | 0.2888 | [0.27, 0.31] |
| 2006 | 5.4739 | [5.19, 5.76] | 0.3910 | [0.37, 0.41] |
| 2007 | 7.4272 | [7.20, 7.66] | 0.5305 | [0.51, 0.55] |
| 2008 | 3.0005 | [2.69, 3.34] | 0.2142 | [0.19, 0.24] |
| 2009 | 2.4799 | [2.27, 2.70] | 0.1771 | [0.16, 0.19] |

**TABLE IV:** *R*_0_ values of Veracruz State with an infection period of 6 days.

| Year | $R_0$ | 95% confidence | $\beta$ | 95% confidence |
| --- | --- | --- | --- | --- |
| 2004 | 6.7127 | [6.26, 7.21] | 1.1187 | [1.04, 1.20] |
| 2005 | 7.7616 | [7.41, 8.10] | 1.2936 | [1.24, 1.35] |
| 2006 | 7.4078 | [7.24, 7.58] | 1.2346 | [1.21, 1.26] |
| 2007 | — | — | — | — |
| 2008 | 2.2133 | [2.02, 2.42] | 0.3688 | [0.34, 0.40] |
| 2009 | 4.64 | [4.37, 4.92] | 0.7740 | [0.73, 0.82] |

**TABLE V:** *R*_0_ values of Veracruz State with an infection period of 14 days.

| Year | $R_0$ | 95% confidence | $\beta$ | 95% confidence |
| --- | --- | --- | --- | --- |
| 2004 | 6.7127 | [6.26, 7.21] | 0.4795 | [0.43, 0.52] |
| 2005 | 7.7616 | [7.41, 8.10] | 0.5544 | [0.53, 0.58] |
| 2006 | 7.4078 | [7.24, 7.58] | 0.5291 | [0.51, 0.54] |
| 2007 | — | — | — | — |
| 2008 | 2.2133 | [2.02, 2.42] | 0.1581 | [0.14, 0.17] |
| 2009 | 4.64 | [4.37, 4.92] | 0.3317 | [0.31, 0.35] |

**TABLE VI:** *R*_0_ values of Guerrero State with an infection period of 6 days.

| Year | $R_0$ | 95% confidence | $\beta$ | 95% confidence |
| --- | --- | --- | --- | --- |
| 2004 | 1.9000 | [1.77, 2.03] | 0.3166 | [0.30, 0.34] |
| 2005 | 5.1573 | [4.93, 5.39] | 0.8596 | [0.82, 0.91] |
| 2006 | — | — | — | — |
| 2007 | 5.1745 | [4.95, 5.40] | 0.8624 | [0.82, 0.90] |
| 2008 | — | — | — | — |
| 2009 | 4.3526 | [3.80, 4.95] | 0.7254 | [0.63, 0.82] |

**TABLE VII:** *R*_0_ values of Guerrero State with an infection period of 14 days.

| Year | $R_0$ | 95% confidence | $\beta$ | 95% confidence |
| --- | --- | --- | --- | --- |
| 2004 | 1.9000 | [1.77, 2.03] | 0.1357 | [0.12, 0.15] |
| 2005 | 5.1573 | [4.93, 5.39] | 0.3684 | [0.35, 0.39] |
| 2006 | — | — | — | — |
| 2007 | 5.1745 | [4.95, 5.40] | 0.3696 | [0.35, 0.38] |
| 2008 | — | — | — | — |
| 2009 | 4.3526 | [3.80, 4.95] | 0.3109 | [0.27, 0.35] |

**TABLE VIII:**
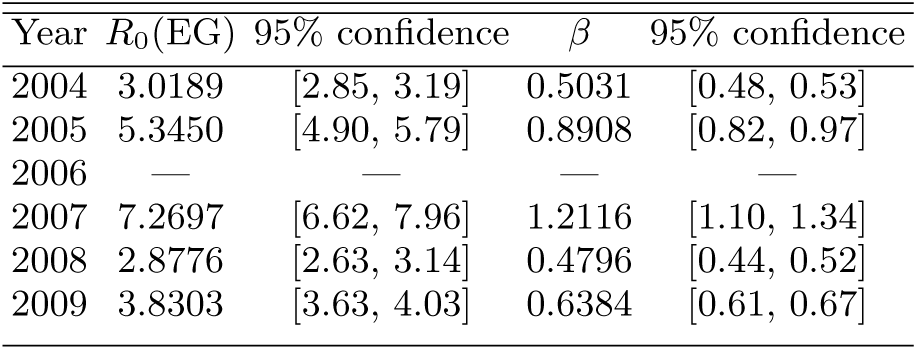
*R*_0_ values of Chiapas State with an infection period of 6 days.

**TABLE IX:** *R*_0_ values of Chiapas State with an infection period of 14 days.

| Year | $R_0$(EG) | 95% confidence | $\beta$ | 95% confidence |
| --- | --- | --- | --- | --- |
| 2004 | 3.0189 | [2.85, 3.19] | 0.2156 | [0.20, 0.23] |
| 2005 | 5.3450 | [4.90, 5.79] | 0.3817 | [0.35, 0.41] |
| 2006 | — | — | — | — |
| 2007 | 7.2697 | [6.62, 7.96] | 0.5192 | [0.47, 0.57] |
| 2008 | 2.8776 | [2.63, 3.14] | 0.2055 | [0.18, 0.22] |
| 2009 | 3.8303 | [3.63, 4.03] | 0.2739 | [0.25, 0.29] |

Finally Table X shows *β*_*i*_ used in the simulations. The starred values are within the confidence interval for *β*_*i*_ (*p-value* ≤ 0.05). On the other hand, those outside of the confidence interval were calculated as “beta optimal values” to approximate the data. Note that, although outside of the confidence interval some of them are relatively close to it.

**TABLE X:**
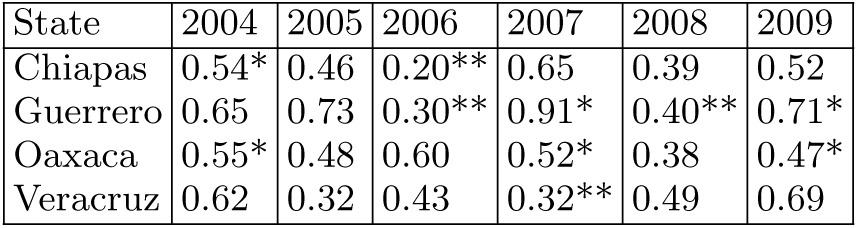
Infection rates *β*_*i*_ for the model simulations.

| State | 2004 | 2005 | 2006 | 2007 | 2008 | 2009 |
| --- | --- | --- | --- | --- | --- | --- |
| Chiapas | 0.54* | 0.46 | 0.20** | 0.65 | 0.39 | 0.52 |
| Guerrero | 0.65 | 0.73 | 0.30** | 0.91* | 0.40** | 0.71* |
| Oaxaca | 0.55* | 0.48 | 0.60 | 0.52* | 0.38 | 0.47* |
| Veracruz | 0.62 | 0.32 | 0.43 | 0.32** | 0.49 | 0.69 |

## References

[1] Angulo, M.T and Velasco-Hernandez Jorge X. Robust qualitative estimation of time-varying contact rates under uncertainty. Epidemics, https://doi.org/10.1016/j.epidem.2018.03.001, 2018.

[2] Burt FJ, Rolph MS, Rulli NE, Mahalingam S, Heise MT (2012) Chikungunya: a re-emerging virus. Lancet 379(9816):662–671. https://doi.org/10.1016/S0140-6736(11)60281X

[3] Cauchemez S, Ledrans M, Poletto C, Quenel P, de Valk H, Colizza V, Boelle PY (2014) Local and regional spread of Chikungunya fever in the Americas. Euro Surveill 19(28).

[4] Carrillo-Valenzo, E., Rogelio Danis-Lozano, Jorge X. Velasco-Hernandez, Gilma Sanchez-Burgos, Celia Alpuche, Irma Lopez, Claudia Rosales, Cecile Baronti Xavier de Lamballerie, Edward C. Holmes and Jose Ramos-Castaneda. Evolution of Dengue Virus in Mco is characterized by frequent lineage replacement. Archives of Virology 155 (9): 1401–1412, 2010.

[5] Cazelles B, Chávez M, Mc Michael AJ, Hales S (2005) Nonstationary Influence of El Ninño on the Synchronous Dengue Epidemics in Thailand. Plos Medicine 2(4):313–3318.

[6] Chakrabarti D, Leskovec J, Faloutsos D, Madden S, Guestrin C, Faloutsos M (2007) Information Survival Threshold in Sensor and P2P Networks. In INFOCOM 2007 26th IEEE International Conference on Computer Communications. IEEE. https://doi.org/10.1109/INFCOM.2007.156

[7] Chakrabarti D, Wang Y, Wang C, Leskovec J, Faloutsos C (2008) Epidemic thresholds in real networks. ACM Trans. Inform. Syst. Secur. 10(4):1–26. https://doi.acm.org/ 10.1145/1284680.128468

[8] Chen J, Beier JC, Cantrell RS, Cosner C, Fuller DO, Guan Y, Zhang G, Ruan S (2018) Modeling the Importation and Local Transmission of Vector-Borne Diseases in Florida: The Case of Zika Outbreak in 2016. Journal of Theoretical Biology 455:342–356. https://doi.org/10.1016/j.jtbi.2018.07.026

[9] Conagua https://www.gob.mx/conagua/acciones-y-programas/sistema-nacional-de-informacion-del-agua-sina.

[10] Cosner C (2015) Models for the effects of host movement in vector-borne disease systems. Mathematical Biosciences 270:192–197. https://doi.org/10.1016/j.mbs.2015.06.015

[11] Datatur https://www.datatur.sectur.gob.mx/SitePages/InfTurxEdo.aspx

[12] Ebi KL, Nealon J (2016) Dengue in a changing climate. Environmental Research 151:115–123. https://doi.org/10.1016/j.envres.2016.07.026

[13] Falcón-Lezama JA, Santos-Luna R, Roman-Pérez S, Martínez-Vega RA, Herrera-Valdez MA, Kuri-Morales AF, et al. (2017) Analysis of spatial mobility in subjects from a Dengue endemic urban locality in Morelos State, Mexico. PLoS One 12(2). https://doi.org/10.1371/journal.pone.0172313

[14] Feng Z, Velasco-Hernández J (1997) Competitive exclusion in a vector-host model for the dengue fever. Journal of Mathematical Biology 35(5):523–544. https://doi.org/10.1007/s002850050064

[15] Gómez S, Arenas A, Borge-Holthoefer J, Meloni S, Moreno Y (2010) Discrete-time Markov chain approach to contact-bases disease spreading in complex networks. Europhys. Lett. 89(3):1–6. https://doi.org/10.1209/0295-5075/89/38009

[16] Gómez S, Arenas A, Borge-Holthoefer J, Meloni S, Moreno Y (2011) Probabilistic framework for epidemic spreading in complex networks. Int. J. Complex Syst. Sci. 1:47–54.

[17] González-Morales N., Núñez-López M, Jose Ramos-Castaneda and Jorge X. Velasco-Hernandez. (2017) Transmission dynamics of two dengue serotypes with vaccination scenarios. Mathematical Biosciences 287:54–71. https://doi.org/10.1016/j.mbs.2016.10.001

[18] Heesterbeek H, Amnderson RM, Andreasen V et al (2015) Modeling infectious disease dynamics in the complex landscape of global health. Science 347 (6227). https://doi.org/10.1126/science.aaa4339.

[19] Khan K, Bogoch I, Brownstein JS, Miniota J, Nicolucci A, Hu W, Nsoesie EO, Cetron M, Creatore MI, German M, Wilder-Smith A (2014) Assessing the Origin of and Potential for International Spread of Chikungunya Virus from the Caribbean. PLOS Currents Out-breaks 6. https://doi.org/10.1371/currents.outbreaks.2134a0a7bf37fd8d388181539fea2da5.

[20] Keymer JE, Marquet PA, Velasco-Hernández JX, Levin SA (2000) Extinction Thresholds and Metapopulation Persistence in Dynamic Landscapes. The American Naturalist 156(5): 478–494.

[21] Leparc-Goffart I, Nougairede A, Cassadou S, Prat C, De Lamballerie X (2014) Chikungunya in the Americas. Lancet 383(9916):514. https://doi.org/10.1016/S0140-6736(14)60185-9.

[22] Levins R (1969) Some demographic and genetic consequences of environmental hetereogeneity for biological control. Bulletin of the EntomologicaI Society of America 15:237–240.

[23] Levins R (1970) Extinction in Some mathematical problems in biology. Mathematical Society of America, Providence, RI.

[24] Liu-Helmersson J, Stenlund H, Wilder-Smith A, Rocklov J (2014) Vectorial Capacity of Aedes aegypti: Effects of Temperature and Implications for Global Dengue Epidemic Potential. PLoS ONE 9(3):e89783. https://doi.org/10.1371/journal.pone.0089783

[25] Fernández López L, Amaku M, Bezerra Coutinho FA, Quam M, Nascimento Burattini M, Struchiner CJ, Wilder-Smith A, Massad E (2016) Modeling Importations and Exportations of Infectious Diseases via Travelers. Bull Math Biol 78(2):185–209. https://doi.org/10.1007/s11538-015-0135-z.

[26] Luz PM, Struchiner CJ, Galvani AP (2010) Modeling Transmission Dynamics and Control of Vector-Borne Neglected Tropical Diseases. PLoS Negl Trop Dis 4(10):e761. https://doi.org/10.1371/journal.pntd.0000761

[27] Madeiros LCC, Castilho CAR, et al. (2011) Modeling the dynamic transmission of Dengue fever: investigating disease persistence. PLoS Negl Trop Dis 5(1):e942, 2011. https://doi.org/10.1371/journal.pntd.0000942

[28] Manore CA et al. (2014) Comparing dengue and Chikungunya emergence and endemic transmission in A.aegypti and A.albopictus. Journal of Theoretical Biology 356:174–191. https://doi.org/10.1016/j.jtbi.2014.04.033.

[29] Marquet P (2002) Metapopulations. In Encyclopedia of Global Environmental Change 2. The Earth system: biological and ecological dimensions of global environmental change. John Wiley and Sons, Chichester.

[30] Martínez-Vega RA, Danis-Lozano R, Díaz-Quijano FA, Velasco-Hernández JX, Santos-Luna R, Román-Pérez S, et al. (2015) Peridomestic Infection as a Determining Factor of Dengue Transmission. PLoS Negl Trop Dis 9(12):e0004296. https://doi.org/10.1371/journal.pntd.0004296

[31] Meloni S, Arenas A, Gómez S, Borge-Holthoefer, J, Moreno Y (2012) Modeling Epidemic Spreading in Complex Networks: Concurrency and Traffic. In Handbook of Optimization in Complex Networks, Springer-Verlag:New York, USA, ISBN 978-1-4614-0753-9.

[32] Obadia T, Haneef R, Pierre-Yves BoÃlle (2012) The R0 package: a toolbox to estimate reproduction numbers for epidemic outbreaks. BMC Medical Informatics and Decision Making 12:147. https://doi.org/10.1186/1472-6947-12-147

[33] Parham, Paul, Waldock, Joanna, Christophides, George K, Hemming, Deborah, Agusto, Folashade, Evans, Katherine J, Fefferman, Nina, Gaff, Holly, Gumel, Abba, Ladeau, Shannon, Lenhart, Suzanne, Mick-ens Ronald E, Naumova, Elena N, Ostfeld, Richard S, Ready, Paul D, Thomas, Matthew B, Velasco-Hernandez, Jorge, Michael, Edwin (2015) Climate, environmental and socio-economic change: weighing up the balance in vector-borne disease transmission. Phil. Trans. R. Soc. B 370(1665):20130551. https://doi.org/10.1098/rstb.2013.0551

[34] San Martin JL, Brathwaite-Dick O (2007) La estrategia de gestión integrada para la prevención y el control del Dengue en la región de las Américas. Rev Panam Salud Publica/Pan Am J Public Health 21(1).

[35] Simmons CP, Ph.D., Farrar J, et al. (2012) Dengue. The New England Journal of Medicine 366(15):1423–1432.

[36] Tatem AJ, Rogers DJ, SI Hay (2006) Global transport networks and infectious disease spread. Advances in Parasitology 62:293–343. https://doi.org/10.1016/S0065-308X(05)62009-X

[37] Undurraga EA, Betancourt-Cravioto M, Ramos-Castañeda J, Martínez-Vega R, Mendez-Galvan J, Gubler DJ, et al. (2015) Economic and Disease Burden of Dengue in Mexico. PLoS Negl Trop Dis 9(3):e0003547. https://doi.org/10.1371/journal.pntd

[38] Wang Y, Chakrabarti D, Wang C, Faloutsos C (2003) Epidemic spreading in real networks: An eigenvalue viewpoint. IEEE SRDS 25–34.

[39] Wesolowski A, Qureshi T, Boni MF et al. (2015) Impact of human mobility on the emergence of dengue epidemics in Pakistan. PNAS 112(38): 11887–11892. https://doi.org/10.1073/pnas.1504964112

[40] Wesolowski A, Erbach-Schoenberg Ez et al. (2017) Multinational patterns of seasonal asymmetry in human movement influence infectious disease dynamics. Nature communications 8(1):2069. https://doi.org/10.1038/s41467-017-02064-4

[41] WHO Dengue control. http://www.who.int/denguecontrol/disease/en/ Accessed 22 April 2018.

[42] Xiao JP, He JF, Deng AP, Lin HL, et al. (2016) Characterizing a large outbreak of dengue fever in Guangdong Province, China. Infectious Diseases of Poverty 5(44). https://doi.org/10.1186/s40249-016-0131-z

[43] Wallinga J, Lipsitch M (2007) How generation intervals shape the relationship between growth rates and reproductive numbers. Proc. R. Soc. B 274(1609):599–604. https://doi.org/10.1098/rspb.2006.3754

[44] Mood AM, Graybill FA Boes DC (1974) Introduction to the theory of statistics. Singapore: McGraw-Hill.

[45] Velasco-Hernández JX, Nññez-López M, Comas-García A, Cherpitel DEN, Ocampo MC (2015) Superinfection between Influenza and RSV Alternating Patterns in San Luis PotosAÃ State, México. PLoS ONE 10(3):e0115674. https://doi.org/10.1371/journal.pone.0115674.

